# Autophagy-Mediated Downregulation of AXL and TIM-1 Promotes Sustained Zika Virus Infection

**DOI:** 10.1101/2024.12.31.630961

**Authors:** Jingyou Yu, Yi-Min Zheng, Megan A. Sheridan, Toshihiko Ezashi, R Michael Roberts, Shan-Lu Liu

**Affiliations:** Center for Retrovirus Research, The Ohio State University, Columbus, OH 43210, USA; Department of Veterinary Biosciences, The Ohio State University, Columbus, OH 43210, USA; Department of Biochemistry, University of Missouri, Columbia, MO 65211; Bond Life Sciences Center, University of Missouri, Columbia, MO 65211; Division of Animal Sciences, College of Agriculture, Food, & Natural Resources, University of Missouri, Columbia, MO 65211; Department of Microbial Infection and Immunity, The Ohio State University, Columbus, OH 43210, USA; Viruses and Emerging Pathogens Program, Infectious Diseases Institute, The Ohio State University, Columbus, OH 43210, USA

**Keywords:** ZIKV, AXL, TIM-1, Autophagy, Down-Regulation

## Abstract

Zika virus (ZIKV) infection can lead to a variety of clinical outcomes, including severe congenital abnormalities. The phosphatidylserine (PS) receptors AXL and TIM-1 are recognized as critical entry factors for ZIKV *in vitro*. However, it remains unclear if and how ZIKV regulates these receptors during infection. In this study, we investigated AXL and TIM-1 expression in human alveolar basal epithelial A549 cells, glioblastoma U87 cells, and embryonic stem cells-derived trophoblast following ZIKV infection. We found that both the Asian strain FSS13025 and the African strain MR766 of ZIKV downregulate AXL, with a milder effect on TIM-1. We identified several ZIKV proteins, notably envelope (E), NS2A, NS3, and NS4B, that contribute to this downregulation. Notably, treatment with lysosomal inhibitor NH_4_Cl or the autophagy inhibitor 3-Methyladenine (3-MA) mitigated the AXL/TIM-1 downregulation, indicating autophagy’s involvement in the process. Importantly, this downregulation facilitates sustained viral replication and promotes viral spread by preventing superinfection and limiting cell death, which is also associated with impaired innate immune signaling. Our findings uncover a mechanism by which ZIKV downregulates entry factors to enhance prolonged viral replication and spread.

**AUTHOR SUMMARY:** Zika virus (ZIKV) infection has been associated with severe birth defects, yet the mechanisms underlying its pathogenesis remain poorly understood. In this study, we investigated phosphatidylserine (PS) receptors AXL and TIM-1 and discovered that they promote ZIKV entry but are downregulated by the virus infection. We identified several ZIKV proteins involved in AXL and TIM-1 down-regulation through an autophagy-mediated process. Mechanistically, this loss of surface receptors protects host cells from superinfection and cell death, while dampening the innate immune response, ultimately promoting viral spread. Our results contribute to a better understanding of ZIKV’s interactions with host cells and offer insight into viral entry, innate signaling, and pathogenesis.

## INTRODUCTION

The Zika virus (ZIKV) outbreaks, particularly across the Americas, have caused substantial public health concerns, especially due to their association with severe congenital defects like microcephaly (1–3). While ZIKV infections are often asymptomatic or mild in adults, the virus poses a substantial risk to fetuses by disrupting the development of neuronal progenitor cells, leading to neurological impairments (4–7) The diverse outcomes of ZIKV infection highlight its complex interactions with host cells, but the mechanisms by which the virus exploits cellular entry factors remain incompletely understood.

Phosphatidylserine (PS) receptors, particularly the TAM family (Tyro3, Axl, Mer) and TIM (T-cell immunoglobulin and mucin domain) family proteins, have emerged as crucial entry factors for many viruses, including ZIKV(11). These receptors facilitate viral entry by interacting with exposed PS on the viral envelope. While studies have highlighted the critical role of AXL and TIM-1 in ZIKV infection, conflicting evidence from *in vivo* models has called into question their necessity for viral pathogenesis (12–14). For instance, AXL-knockout mice exhibit ZIKV susceptibility similar to wildtype controls (12), suggesting that receptor usage may vary by tissue or cellular context. PS normally resides on the inner leaflet of the plasma membrane but can be translocated to the outer surface during specific physiological or pathological processes, such as blood-clotting, sperm maturation or viral infection (15–19). PS exposure is considered as a hallmark of early apoptosis, and PS receptors on phagocytic cells recognize the exposed PS, leading to the clearance of apoptotic cells thus preventing local inflammation (20, 21). Importantly, many viruses have evolved to exploit this “apoptotic mimicry” by incorporating PS into their envelope (22–24), thereby enhancing their ability to bind and enter host cells via PS receptors. This has led to the proposal of PS receptors, including AXL and TIM-1, as candidate receptors for several viruses, such as HAV (25), DENV (26), EBOV (27), and others (28, 29). Adding to the debate over the specificity of PS receptors for viruses, recent studies have highlighted their critical roles in ZIKV infection and pathogenesis, especially AXL and TIM-1 (5, 6, 11). AXL is notably abundant in immune-privileged sites, such as neurons, trophoblasts, and testes, where ZIKV robustly replicates. AXL binding to PS requires an adaptor molecule, such as PROS1 or GAS6, whereas TIM-1 can directly bind PS with high affinity (28).

Superinfection interference, where a prior viral infection prevents subsequent infection with the same or related virus, has been observed for numerous viruses (30). This phenomenon involves mechanisms such as receptor site competition, RNA replication interference, and virus budding (31, 32). Viruses can downregulate receptors through receptor-mediated endocytosis, leading to lysosomal degradation of viral entry factors. This strategy benefits the virus by preventing the accumulation of viral receptors or cofactors that might trigger innate immune responses, reducing receptor-mediated signaling that could lead to cell death, and/or preventing viral recombination due to excessive viral load. While superinfection interference mechanisms are well-documented in other viruses including HIV (33) and West Nile virus (34), the possible role of PS receptor modulation in ZIKV infection remains unknown.

In this study, we investigate the regulatory effects of ZIKV infection on the expression of AXL and TIM-1 in various cell types. We aim to elucidate the mechanisms through which ZIKV modulates these receptors and the potential implications for viral replication, superinfection, and immune evasion.

## RESULTS

### CRISPR knockout of human AXL and TIM-1 differentially reduces ZIKV entry

PS receptors are crucial cellular factors that facilitate the entry of various viruses, including retroviruses (19), filoviruses (27, 35), flaviviruses (26, 36) and others (37). To explore the roles of two key PS receptors, AXL and TIM-1, in modulating ZIKV infection, we established A549 CRISPR/Cas9 knockout cell lines to deplete AXL or TIM-1 expression, designated as AXL KO and TIM-1 KO, respectively. The expression levels of AXL and TIM-1, in both total and the cell surface, were determined by immunoblotting (**Fig. 1A**), flow cytometry staining (**Fig. 1B**), and immunofluorescence imaging (**Fig. S1A**). To minimize cell death caused by ZIKV replication, we infected these KO cells with ZIKV MR766 strain at a relatively low MOI=0.1 for 24 hours, and the subsequent ZIKV infection rates were quantified by staining ZIKV E protein measured by flow cytometry. Compared with Scramble KO cells, where 48.4% were ZIKV E positive, AXL KO and TIM-1 KO showed 30.6% and 18.9% ZIKV E-positive cells, respectively (**Fig. 1C**). The ZIKV nonstructural protein 1 (NS1) accumulations in these cell lines were examined at 0-, 24-, and 36-hour post-infection (PI) by immunoblotting. We found that the ZIKV NS1 expression level was higher in A549 Scramble KO cells compared to AXL KO and TIM-1 KO cells at both 24- and 36-hour PI (**Fig. 1D**). The NS1 expression in TIM-1 KO was almost undetectable, indicating an almost complete block of virus infection. Notably, at 36 hours PI, the NS1 accumulation was lower than that at 24 hours in all cell lines, likely due to increased cell death induced by ZIKV infection, as evidenced by the decreased expression of the loading control, β-actin (**Fig. 1D**).

**Figure 1.**
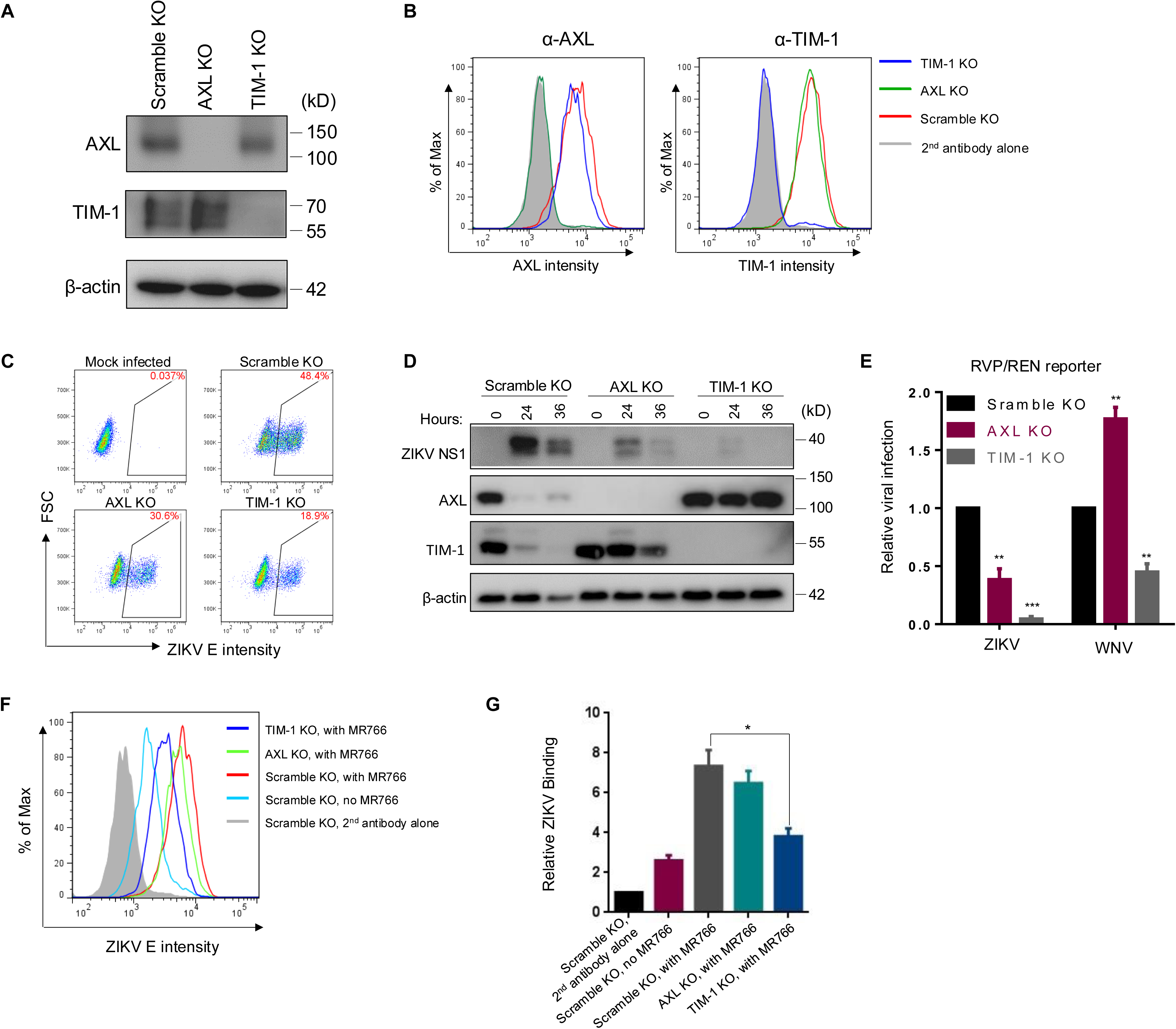
CRISPR knockout of human AXL and TIM-1 differentially modulates ZIKV infection and entry. AXL and TIM-1 expression levels in stable A549 Scramble KO, AXL KO, and TIM-1 KO cells were examined by immunoblotting (**A**) or immunostaining (**B**). (**C**) A549 Scramble KO, AXL KO, and TIM-1 KO cells were infected with MR766 at MOI=1 for 24 hours. Cells were processed for E protein staining (4G2) and analyzed by flow cytometry. Mock-infected cells served as negative controls. The percents of E protein-positive cells were displayed. (**D**) The total protein level of ZIKV NS1, AXL and TIM-1 expression following 0, 24 or 36 hours of infection in A549 Scramble KO, AXL KO and TIM-1 KO cells was determined by western blotting. (**E**) The ZIKV-E or WNV-E pseudotyped one-round RVP/WNV-REN reporter viruses were applied to A549 Scramble KO, AXL KO, and TIM-1 KO cells for 36 hours. Cell lysates were used for measuring the *Renilla* luciferase activity. Data shown are mean ± SD from three independent experiments. Statistical significance was determined by Student’s two-sided t-test, **p < 0.01, ***p < 0.001. (**F-G**) Virus binding. A549 Scramble KO, AXL KO, and TIM-1 KO cells (1 × 10⁶ cells) were mixed with 5 × 10⁶ PFU MR766 virus for 2 hours on ice. After three washes with 1 × PBS, cells were stained with 4G2 and FITC-anti-mouse on ice. The binding efficiency of MR766 on A549 cells was analyzed by flow cytometry and plotted as histograms **(F)**. Mock-infected Scramble KO cells served as a negative control, and second antibody staining alone served as a staining control. Geometric means of ZIKV E staining signals are as follows: second antibody alone: 490; no MR766, 1183; Scramble KO with MR766, 3312; AXL KO with MR766, 2962; TIM-1 KO with MR766, 1735. Quantitative analysis of ZIKV binding to three cell lines from three independent experiments; data are shown as mean ± SD. Statistical significance was determined by Student’s two-sided t-test, **p < 0.01, ***p < 0.001.

It is well established that PS receptors augment viral entry (22). We next used one-round replicon viral-like particles (RVP/REN) to test the ZIKV and West Nile virus (WNV) entry into these three cell lines. As shown in **Fig. 1E**, knockout of AXL and TIM-1 reduced the ZIKV MR766 RVP/REN entry by 3- and 8-fold, respectively. In contrast, the WNV RVP entry was reduced by approximately 2-fold in TIM-1 KO cells, yet surprisingly promoted by 1.7-fold in AXL KO cells, indicating that AXL and TIM-1 differentially modulate flavivirus uptake.

We further investigated if virus binding was affected by AXL and TIM-1 knockout. To this end, we incubated concentrated MR766 viral particles with respective A549 cell lines on ice for 2 hours. Following extensive washes, virions bound to the cell surface were determined by flow cytometry by using an anti-ZIKV E (4G2) antibody, the efficiency of which was quantified by calculating the geometric mean of ZIKV E protein fluorescence signals (**Fig. 1E**). Knockout of AXL and especially TIM-1, strongly reduced the ZIKV binding to A549 cells (**Figs. 1F** and **1G**), suggesting that TIM-1 is more critical for ZIKV entry than AXL, at least in A549 cells.

### ZIKV infection induces AXL downregulation in human epithelial, glioblastoma and trophoblast cells

Given that ZIKV replicates robustly in AXL and TIM-1 expressing A549 cells, we sought to investigate whether and how ZIKV infection may modulate AXL and TIM-1 in this and related cell types. ZIKV MR766 was applied to A549 cells at MOI of 0, 0.05, 0.2, and 1.0 for 24 hours, respectively, during which only mild cytopathic effects were observed (less than 20% OD_450_ difference based on WST-1 assay). ZIKV infection rates were then assessed by intracellular staining for ZIKV E protein. As shown in **Fig. 2A**, the fluorescence intensity of ZIKV E protein increased with a growing MOI from 0 to 1.0 being used (**Fig. 2A**, top left). Interestingly, we found that, concomitant with the increased expression of ZIKV E expression in A549 cells, the AXL expression was gradually decreased, both on the cell surface and at the total protein level (**Fig. 2A**, middle and bottom left). TIM-1 expression exhibited a similar trend of downregulation, albeit to a lesser extent (**Fig. 2A**, middle and bottom right). In contrast to the downregulation of AXL and TIM-1, CD81, another cell surface marker, which was stained in parallel, remained unchanged in A549 cells, even when infected with the highest MOI=1 of ZIKV MR766 used (**Fig. 2A**, top right). These results together suggest that ZIKV infection specifically downregulates AXL and TIM-1, including from the cell surface.

**Figure 2.**
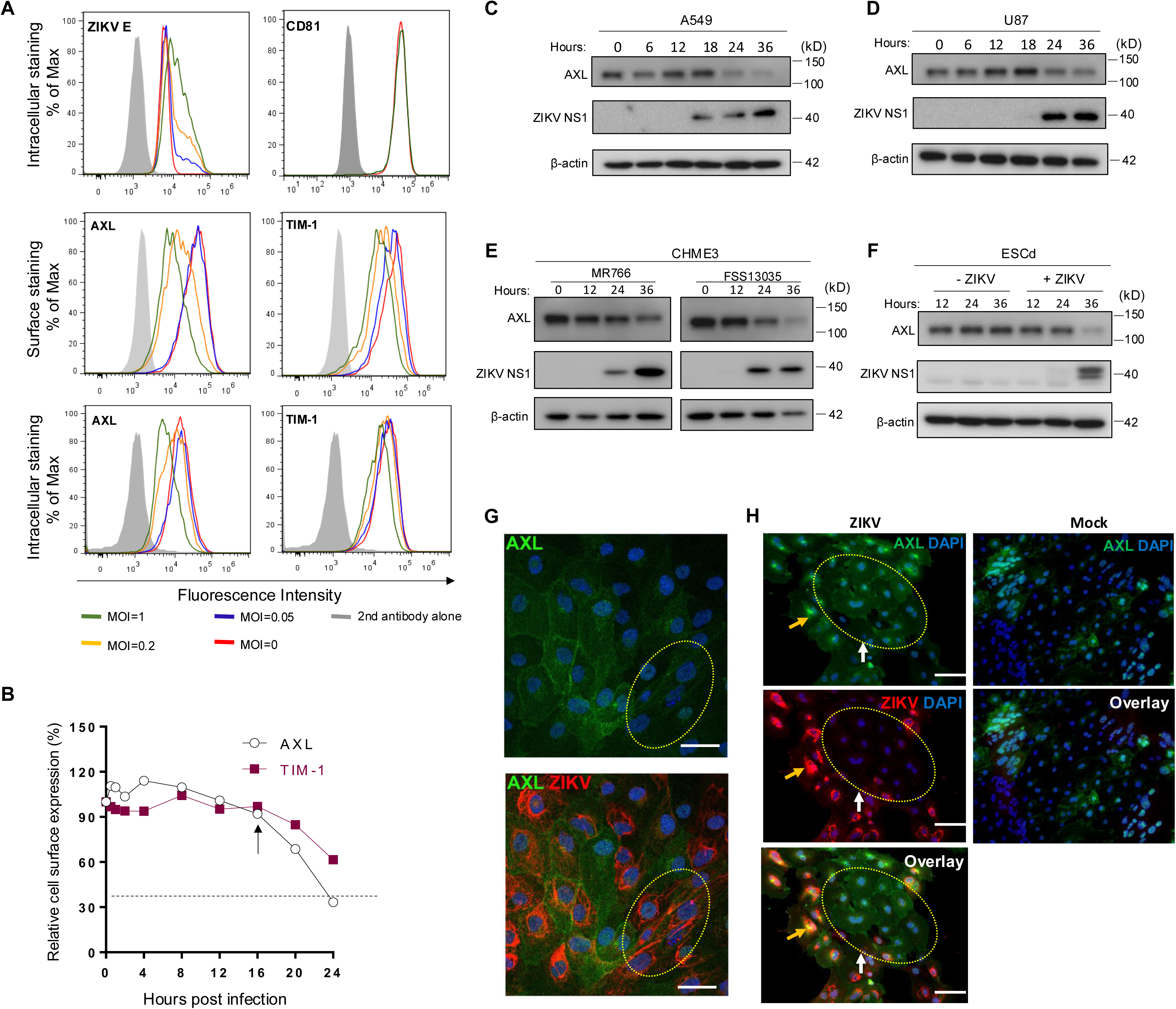
ZIKV infection downregulates AXL in human epithelial, glioblastoma and trophoblast cells. (**A**) A549 cells were infected with ZIKV MR766 at MOI=0, 0.05, 0.2, or 1 for 24 hours. ZIKV E protein was intracellularly stained with an anti-flavivirus E protein antibody (4G2). AXL and TIM-1 proteins were stained to examine both cell surface and intracellular expressions using their specific antibodies. CD81 expression upon ZIKV infection was examined by the cell surface staining. <u>Geometric means of intracellular ZIKV E protein staining</u>: second antibody alone, 552; MOI=0, 4380; MOI=0.05, 5904; MOI=0.2, 9219; MOI=1, 12567. <u>Geometric means of the cell surface AXL protein staining</u>: second antibody alone, 552; MOI=0, 37900; MOI=0.05, 34800; MOI=0.2, 9788; MOI=1, 4710. <u>Geometric means of the cell surface TIM-1 protein staining</u>: second antibody alone, 552; MOI=0, 41300; MOI=0.05, 31900; MOI=0.2, 9788; MOI=1, 8412. Bottom left: <u>Geometric means of intracellular AXL</u>: second antibody alone, 952; MOI=0, 38998; MOI=0.05, 36577; MOI=0.2, 27843; MOI=1, 9876. <u>Geometric means of intracellular TIM-1 protein staining:</u> second antibody alone, 952; MOI=0, 47665; MOI=0.05, 45994; MOI=0.2, 43223; MOI=1, 26757. <u>Geometric means of the cell surface CD81 protein staining</u>: second antibody alone, 889; MOI=0, 29891; MOI=1, 30010. (**B**) Kinetics of AXL and TIM-1 expression on the cell surface upon ZIKV infection within 24 hours. Arrows indicate the approximate time point when downregulation occurred. (**C-F**) ZIKV infection downregulates AXL in A549 cells (**C**), U87 cells (**D**), CHME3 cells (**E**), and trophoblasts differentiated from stem cells (ESCd) (**F**), as examined by immunoblotting. ZIKV African strain MR766 (**C-D**), Asian strain FSS13025 (**E**), or Dakar strain (**F**) were applied for 0, 12, 24, and 36 hours. (**G**) ESCd were infected with ZIKV Dakar strain for 36 hours, and AXL and ZIKV E were detected by immunofluorescence with specific antibodies. Yellow oval indicates area with ZIKV infection. (**H**) Subcellular localizations of AXL in ZIKV-infected ESCd cells were analyzed by immunofluorescence assay. Yellow and white arrows indicate ZIKV positive and negative cells, respectively. Red, ZIKV E; green, AXL; blue, DAPI. Scale bars: 50 μm.

We next examined the dynamics of AXL and TIM-1 downregulation at the cell surface over a 24-hour period following ZIKV infection. As shown in **Fig. 2B**, the downregulation of AXL and TIM-1 began around 16 hours post-ZIKV infection, which largely correlated with the timing of viral protein NS1 expression (**Fig. 2C)**, implying that this downregulation likely resulted from the *de novo* synthesis of ZIKV proteins rather than viral RNA amplification. Immunoblotting analysis of total AXL and TIM-1 protein levels in the lysate of A549 cells confirmed that both proteins began to decrease between 12 or 18- and 24-hours post-infection (**Fig. 2C and Fig. 1D**). To explore if this downregulation is conserved across different cell types, we determined AXL expression in neural glioblastoma U87 (**Fig. 2D**) and microglia CHME3 cells (**Fig. 2E**) upon ZIKV infection. Both African strain MR766 and Asian strain FSS13025 were capable of downregulating AXL expression, despite slightly different kinetics (**Fig. 2D-E**). However, TIM-1 expression was not detected in these neuronal cells, likely due to low endogenous expressions.

In addition to U87 and CHME3 cells, we tested stem cell-derived trophoblast cells, which have been shown to be a primary target of ZIKV (38, 39). Consistent with the results shown above, we observed an increased ZIKV NS1 expression in trophoblasts, which was inversely correlated with a decreased AXL expression (**Fig. 2F**). We further examined AXL expression in trophoblasts by immunofluorescence staining. We found lower levels of AXL expression in areas where an obvious level of ZIKV E was detected (**Fig. 2G and Fig. S2**). Additionally, ZIKV-infected cells exhibited a greatly increased intracellular staining of AXL as compared to its normal cell surface expression, suggesting that ZIKV infection likely sequesters AXL in intracellular compartments and subsequently target it for degradation (**Fig. 2H**). Collectively, these data demonstrate that ZIKV infection results in downregulation of AXL and TIM-1, although the pattern can be cell type dependent.

### ZIKV infection induces autophagy, mediating AXL downregulation

We investigated how ZIKV infection downregulates the expression of AXL and TIM-1. To determine which ZIKV protein(s) are responsible for this downregulation, we co-expressed each of the ten ZIKV proteins, namely E, PrM, C, NS1, NS2B, NS2C, NS3, NS4B, NS4B, and NS5, alongside AXL or TIM-1 in HEK293T cells, and assessed their effects on AXL or TIM-1 expression. As shown in **Fig. 3A**, compared to vector control, co-expression of each of ten ZIKV proteins led to differential patterns of decreases in AXL expression. Among these, ZIKV E, NS2A, NS3, and NS4B showed the most potent effect on downregulating AXL.

**Figure 3.**
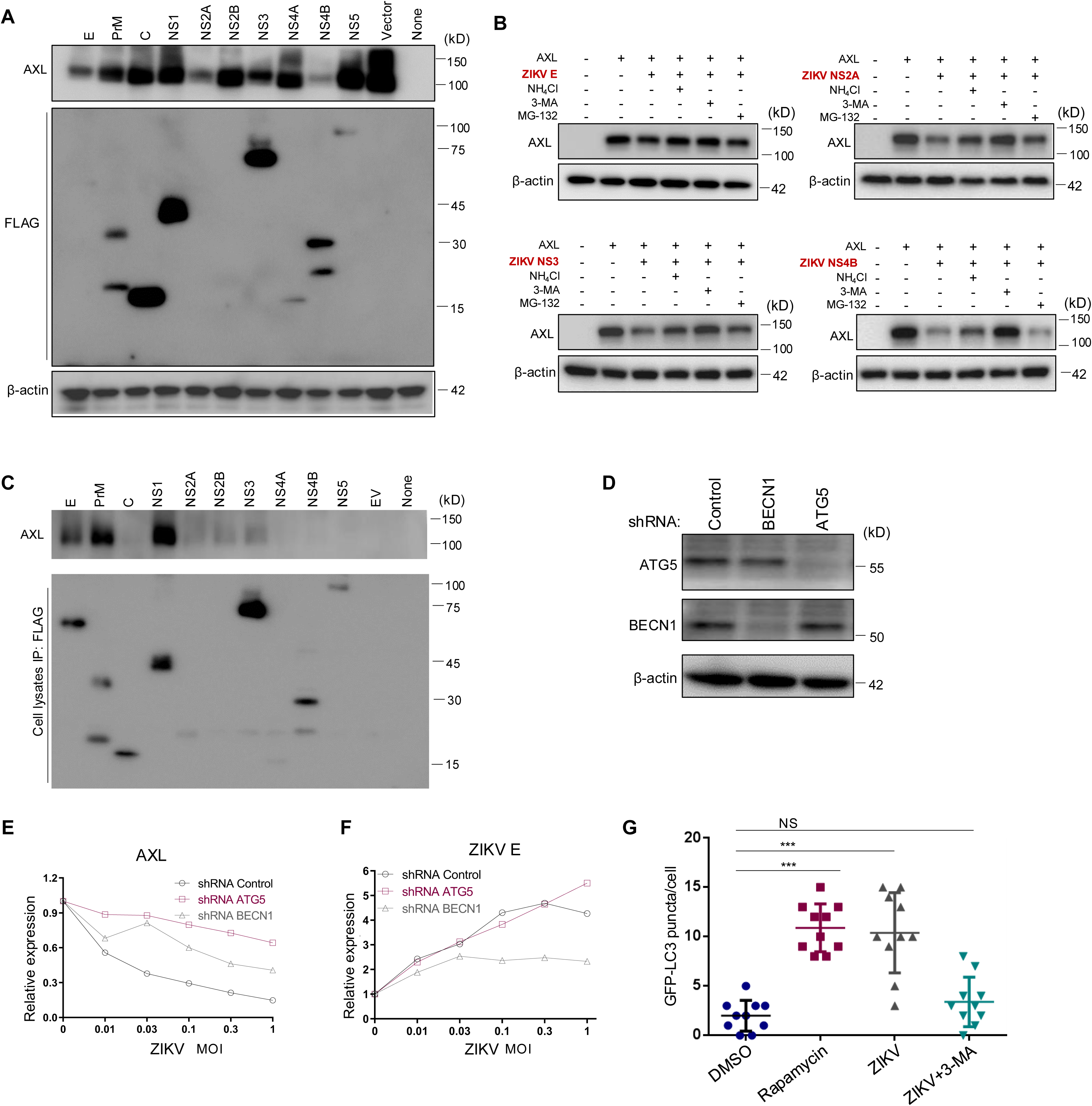
ZIKV infection-induced autophagy mediates AXL downregulation. (**A**) A plasmid expressing AXL was co-transfected along with FLAG-tagged ZIKV constructs into HEK293T cells for 36 hours. Immunoblotting was performed to detect the AXL and ZIKV protein expression using AXL-specific or anti-FLAG antibodies. (**B**) HEK293 cells co-transfected with AXL and ZIKV E, NS2A, NS3, or NS4B were treated with autophagy inhibitor 3-MA, lysosomal inhibitor NH_4_Cl, or proteosomal inhibitor MG-132 for 8 hours. Cell lysates were harvested and processed for immunoblotting of AXL and β-actin as loading control. (**C**) A plasmid expressing AXL was co-transfected with FLAG-tagged ZIKV constructs into HEK293T cells, cells were lysed by RIPA buffer, proteins were pulled down with anti-FLAG-conjugated beads and immunoblotted by using anti-AXL or FLAG antibodies. (**D**) Immunoblotting of Beclin 1 and ATG5 expression in A549 cells expressing shRNA Control, shRNA BECN1, or shRNA ATG5. (**E-F**) A549 shRNA control, shRNA ATG5, or shRNA BECN1 cells were infected with ZIKV MR766 at MOI=0, 0.01, 0.03, 0.1, 0.3, or 1 for 24 hours, and the cell surface AXL (left) or intracellular ZIKV E (right) expression was examined with specific antibodies. The geometric means of fluorescence signals without ZIKV infection was set to 1.0 for comparisons. Data are representatives of two independent experiments with similar results. (**G**) A LC3-GFP plasmid was transfected into A549 cells for 24 hours, followed by treatment with DMSO or Rapamycin for 6 hours, or infection with MR766 for 24 hours with or without 3-MA. The number of GFP-LC3 puncta per cell was quantified by counting 10 typical live cells and plotted. Statistical significance was determined by Student’s two-sided t-test, **p < 0.01, ***p < 0.001.

We next evaluated the possible mechanisms by which these four ZIKV proteins downregulate AXL. To end this end, we applied 20 mM NH_4_Cl to increase lysosomal pH and block protein destruction, 2.5 mM 3-MA to inhibit autophagy, or 20 µM MG-132 to block proteasomal degradation, respectively. As shown in **Fig. 3B**, treatment of cells with 3-MA largely restored the level of AXL downregulated by all four ZIKV proteins, especially NS2B, NS3 and NS4B. NH_4_Cl treatment partially rescued ZIKV E-, NS3- and NS4B-mediated AXL downregulation. In contrast, MG-132 treatment had no obvious effects on AXL downregulation (**Fig. 3B**). These results together suggest that the autophagy/lysosomal degradation pathway is likely involved in the AXL downregulation by ZIKV proteins.

To test if these ZIKV proteins became associated with AXL, thereby leading to its subsequent destruction, we performed co-immunoprecipitation (Co-IP) assays using anti-FLAG beads to pull down each of the 10 FLAG-tagged ZIKV complexes, followed by immunoblotting with an anti-AXL antibody. As shown in **Fig. 3C**, a strong association was observed of AXL with ZIKV E, PrM, and NS1, a weaker interaction with NS2A, NS2B, and NS3, but a limited to no interaction with ZIKV NS2A, NS3 or NS4B. These results suggested that the downregulation of AXL by ZIKV proteins did not strictly correlate with their associations evidenced in the Co-IP assays.

We then examined the role of autophagy in AXL downregulation by ZIKV infection. We first generated A549 stable cell lines after transducing lentiviral shRNA vectors directed against BECN1 and ATG5, two key autophagy-related proteins. As shown in **Fig. 3D**, lentiviral shRNA transduction resulted in an efficient knockdown of BECN1 and ATG5 expression in the A549 cells. We then infected these stably transfected cells with ZIKV MR766 over a range of MOIs, and, 24 hours later, quantified ZIKV E and AXL expression by flow cytometry. We observed that, in response to ZIKV infection, expression of AXL was largely retained in ATG5 knockdown cells although only partially so in BECN1 knockdowns compared to shRNA control cells (**Fig. 3E**), reinforcing the notion that autophagy-mediated degradation likely plays a critical role in AXL downregulation. As expected, increased viral inputs led to an elevated ZIKV E accumulation in A549 cells (**Fig. 3F**). However, a comparable increase in ZIKV E was noted in the ATG5 knockdown cells, while increases were more modest in the BECN1 knockdowns (**Fig. 3F**), was substantially suppressed, suggesting that ZIKV replication could have benefited from the process of autophagy.

We next sought to determine if ZIKV infection itself induces autophagy. To this end, we created an autophagy reporter cell line after transducing A549 cells with a lentivector expressing LC3B fused with GFP at the N-terminus (eGFP-LC3) (40). These cells, which express GFP in response to autophagosome assembly, were treated with either Rapamycin (a classical autophagy inducer, dissolved in DMSO) or with DMSO as a control. Alternatively, cells were infected with an MR766 ZIKV strain alone, in the presence or absence of 3-MA (a PI3K-autophagy inhibitor). Compared to DMSO-treated cells, which exhibited few LC3-GFP puncta, Rapamycin-treated cells showed a marked increase in puncta (**Fig. 3G**). Similar to previous reports (41), we found that ZIKV infection also increased number of LC3 puncta, which were reduced by the 3-MA treatment (**Fig. 3G**). These data collectively suggest that ZIKV infection induces autophagy, which, in turn, contributes to the ZIKV protein-mediated downregulation of AXL.

### Depletion of AXL and TIM-1 attenuates ZIKV infection-induced autophagy and apoptosis

To assess the impact of AXL and TIM-1 knockdown on ZIKV replication and spread, we infected three A549 cell lines (AXL KO, TIM-1 KO cells, and control scramble KO) with either MR766 or the Cambodian strain of ZIKV FSS13025 for 24 hours (MOI = 0.1). We observed a marked attenuation of cytopathic effect (CPE) in AXL KO cells, and to a lesser extent, in TIM-1 KO cells, compared with scramble KO cells (**Fig 4A and Fig. S3A**). Quantitative analysis revealed a significantly increased cell viability in AXL KO cells at 24 hours PI and onward, whereas TIM-1 KO cells showed smaller but significant improvement in viability at 48 hours PI (**Fig. 4B**). These results together suggested that depletion of TIM-1, and especially AXL, can mitigate ZIKV infection-induced cell death.

**Figure 4.**
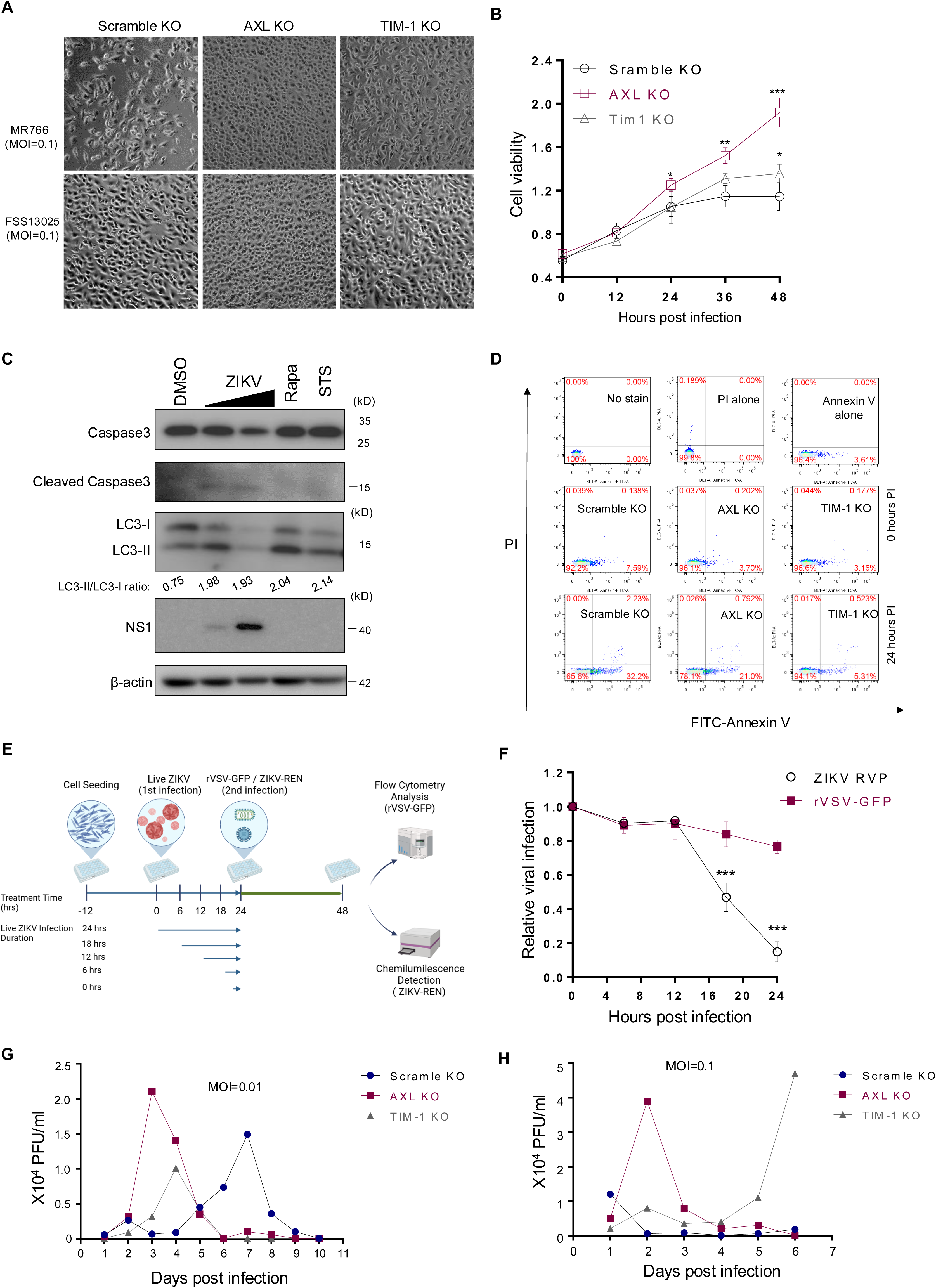
ZIKV replication is sustained in AXL knockout cells, correlating with reduced superinfection and cell death. (**A**) A549 Scramble KO, AXL KO, and TIM-1 KO cells were infected with ZIKV MR766 or FSS13025 at MOI=0.1 for 24 hours. Cells were imaged after floating cell debris was removed. (**B**) A549 Scramble KO, AXL KO, and TIM-1 KO cells were infected with ZIKV and at 0, 12, 24, 36, and 48 hours, the viability was quantified by WST-1 assay. Data shown are mean ± SD from three independent experiments. Statistical significance was determined by Student’s two-sided t-test, *p < 0.05, **p < 0.01, ***p < 0.001. (**C**) A549 cells were treated with DMSO, Rapamycin (Rapa), Stausporin (STS) for 6 hours or infected with MR766 for 24 hours at MOI=0.1 or MOI=1. Cells were lysed and subjected to immunoblotting using anti-Caspase-1, caspase-3, LC3 and ZIKV NS1 specific antibodies; β-actin served as loading control. The intensity of LC3-II and LC-I bands was quantified, and the LC3-II vs. LC3-I ratio was calculated and shown. (**D**) A549 Scramble KO, AXL KO, and TIM-1 KO cells were infected with ZIKV (MOI=0.1) for 0 or 24 hours, and phosphatidylserine (PS) exposure on the cell surface was measured by flow cytometry using with FITC-Annexin V; dead cells were stained with propidium iodide (PI). (**E**) Schematic of two-round ZIKV infection procedures. A549 cells were first infected with MR766 for 0, 6, 12, 18, or 24 hours (MOI=1) and subsequently challenged with either ZIKV-REN RVP or rVSV-GFP. The impact of first-round infection on the second round was determined by measuring the *Renilla* luciferase activity or GFP signal. (**F**) The relative rate of second-round infection was calculated by setting the infection without the first-round infection to zero. Data shown are mean ± SD from three independent experiments. Statistical significance was determined by Student’s two-sided t-test, ***p < 0.001. (**G-H**) Long-term replication of MR766 in three A549 cell lines was performed at MOI=0.001 or 0.01. Supernatants containing viruses were collected daily until cell death occurred, and virus titers at each time point were determined by plaque assay.

We next sought to investigate the cell death pathways that ZIKV infection might activate, as well as how AXL and TIM-1 downregulation might influence these signaling events. To this end, we performed immunoblots of proteins of interest involved in apoptosis and autophagy in cells following ZIKV infection. Similar to Rapamycin treatment, ZIKV infection increased the conversion of LC3-I to LC3-II, i.e., LC3-II/LC3-I ratio, compared to the DMSO-treated control at a low MOI. However, a high MOI of infection dramatically reduced total LC3 expression (**Fig. 4C**), making the calculation of LC3-II/LC3-I ratio difficult. Nonetheless, these data indicated that ZIKV infection indeed triggers autophagy as observed in some previous reports (42). Of note, ZIKV infection also caused an increased level of cleaved CASP3 (caspase-3), an indicator of apoptosis induction (**Fig. 4C**), suggesting that ZIKV infection can trigger both autophagy and apoptosis, perhaps collectively contributing to the observed cell death.

To directly measure the impact of TIM-1 and AXL KO on cell death induced by ZIKV infection, we performed ANXA5 (annexin V) staining to detect exposed phosphatidylserine (PS), an early marker of apoptosis, as well as propidium iodide (PI) staining to mark dead cells. As shown in **Fig. 4D**, in the absence of ZIKV infection (MOI=0.1), i.e., 0 hours post-infection, AXL KO and TIM-1 KO cells exhibited lower levels of ANXA5-positive staining, i.e., 3.70% and 3.16%, respectively. This was in comparison to Scramble KO cells, where ANXA5-stained cells numbered 7.59%, suggesting that knockout of these PS receptors reduces the basal level of PS flipping. These results were consistent with our previous findings that expression of PS receptors, especially TIM proteins, induces PS flipping (43). At 24-hour PI, ANXA5-positive cells were increased to 32.2% in scramble KO cells in contrast to the 21.0% and 5.31% detection in AXL KO and TIM-1 KO cells, respectively. These results together demonstrated that depletion of TIM-1, and to a lesser extent AXL, reduced PS flipping.

### Depletion of AXL and TIM-1 limits ZIKV superinfection and promotes virus spread

Prevention of superinfection is known to benefit virus replication (30). To test whether ablation of AXL and/or TIM-1 would have any impact on superinfection and virus spread, we carried out a short-term infection of A549 cells with a high MOI = 1 of ZIKV MR766 for a period of 0, 6, 12, 18, and 24 hours, respectively to allow a strong downregulation of AXL and TIM-1.

These treatments were followed by infection of the cells either with one-round reporter virus RVP/ZIKV-Luc, or a replication-competent rVSV-G-GFP, which served as a control, for another 24 hours. The second round of ZIKV and VSV infection was quantified by measuring *Renilla* luciferase activity and GFP fluorescence by flow cytometry, respectively. As shown in **Figs. 4E** and **4F**, the second round of ZIKV infection (entry) was reduced after 12 h of first-round of ZIKV infection, correlating with the kinetics of AXL and TIM-1 downregulation (**Fig. 2B**). In contrast, the rVSV-G-GFP infection was not substantially affected (**Fig. 4F)**.

We next performed a long-term replication study of up to 10 days with infectious ZIKV MR766 in A549 Scramble KO, AXL KO, and TIM-1 KO cells using a low MOI=0.01. We found that ZIKV replication in Scramble KO A549 cells peaked at day 7, which was much delayed to compared to virus replication in AXL KO and TIM-1 KO cells, which peaked at 3 and 4 days PI, respectively (**Fig. 4G**). Notably, when a higher MOI=0.1 was used, ZIKV replication peaked in Scramble KO A549 cells at day 1, with approximately 1.0 × 10^4^ plaque-forming units (PFUs) virions per ml produced. This was in contrast to the amount of 4.0 × 10^4^ and 5.0 × 10^4^ PFU virions produced per ml, which peaked at day 2 and day 6 PI in AXL KO and TIM-1 KO cells, respectively (**Fig. 4H**). Taken together, these results suggested that knockdown of both TIM-1 and AXL promotes prolonged ZIKV replication, although the initial phase of viral entry is impaired because of the lack of these critical host factors for particle binding.

### AXL-deficiency dampens the type I IFN response and inflammatory signaling

The innate immune response is critical for restricting viral infection. To explore the possible roles of AXL and TIM-1 in modulating type I interferon (IFN) and inflammatory signaling, we infected A549 scramble KO, AXL KO, and TIM-1 KO cells with a high MOI (MOI = 5) in order to minimize the defect of AXL and TIM-1 on ZIKV entry, thus permitting a relatively comparable level of incoming ZIKV genomic RNA to trigger subsequent signaling cascades. Indeed, 18 hours after infection, AXL KO only showed a modest ∼20% reduction of ZIKV C mRNA expression while TIM-1 KO exhibited a greater (approximately 50%) reduction of incoming viral RNAs (**Fig. 5A**).

**Figure 5.**
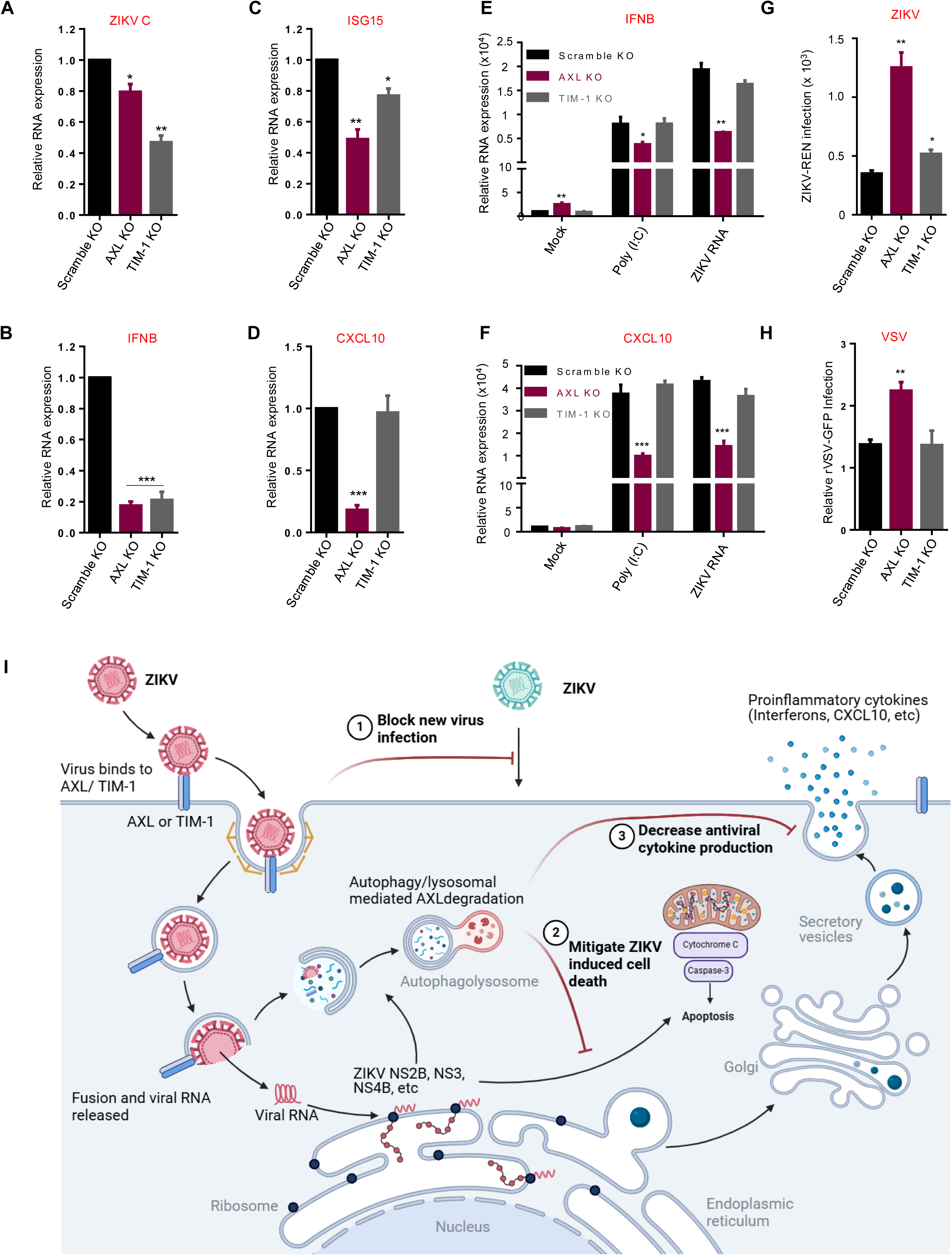
AXL deficiency dampens antiviral type I IFN response and inflammatory signaling, which in turn promotes ZIKV infection. A549 Scramble KO, AXL KO, and TIM-1 KO cells were infected with ZIKV at an MOI of 5. Eighteen hours after infection, total cellular RNA was extracted and the mRNA levels of ZIKV C (**A**), IFNB (**B**), ISG15 (**C**), and CXCL10 (**D**) were quantified by qRT-PCR. Alternatively, Poly (I:C) or ZIKV genomic RNA (vRNA) was transfected into A549 Scramble KO, AXL KO, and TIM-1 KO cells, and 18 hours after transfection, mRNA levels of IFNB (**E**) and CXCL10 (**F**) were measured by qRT-PCR. Mock-treated cells served as negative controls. (**G-H)** Conditioned media collected from the transfected A549 cells were added to Vero cells for 24 hours. Subsequently, ZIKV-REN or rVSV-GFP reporter viruses were applied to the treated cells for 24 hours. Cells were then lysed for *renilla* luciferase activity (**G**) or GFP (**H**). In all cases, data are shown as mean ± SD from three independent experiments. Statistical significance was determined by a two-sided Student’s t-test, *p < 0.05, **p < 0.01, ***p < 0.001. (**I**) Working model for ZIKV infection-induced AXL or TIM-1 downregulation by the autophagy/lysosomal degradation pathway. See text in the Discussion section.

We then measured the expression level of IFNB and ISG15, two key markers of type I interferon signaling, as well as the pro-inflammatory cytokine CXCL10. As shown in **Figs. 5B** and **5C**, both IFNB and ISG15 were significantly lower in AXL and TIM-1 KO cells compared to scramble KO cells. Notably, CXCL10 expression was reduced by ∼80% in AXL KO cells, but not in TIM-1 KO cells (**Fig. 5D**). Together, these data suggest that AXL potentiates the type I IFN and pro-inflammatory signaling, and that the observed effect of TIM-1 KO was likely due to the reduced level of ZIKV RNA input at entry.

To expand above findings, we transfected double-stranded RNA mimic poly (I:C) and ZIKV-Rluc viral genomic RNA separately into these A549 cell lines and measured the expression of IFNB and CXCL10. We observed decreased production of both IFNB (**Fig. 5E**) and CXCL10 (**Fig. 5F**) in AXL KO but not in TIM-1 KO cells following transfection, confirming that depletion of AXL in A549 cells attenuates RNA ligand-induced innate immune signaling.

To assess the functional impact of altered IFNB and CXCL10 expression on viral infection, we transfected ZIKV-RLuc genomic RNA into A549 scramble KO, AXL KO, or TIM-1 KO cells for 24 hours, thus allowing the induction of innate immune effectors, and harvested the cell supernatant containing ZIKV virions. After UV-inactivating the newly produced ZIKV viruses, the conditioned medium was applied to Vero cells for 12 hours before being infected with either ZIKV-RLuc or rVSV-GFP viruses. As shown in **Figs 5G** and **5H**, both ZIKV and VSV replicated more rapidly in Vero cells cultured in AXL KO cell media compared to media from scramble KO and TIM-1 KO cells, implying that less antiviral effectors were produced in AXL KO cells. Overall, these data suggested that depletion of AXL results in impaired antiviral innate immune responses, thus contributing to enhanced viral infection and spread.

## DISCUSSION

This study uncovers the molecular mechanism by which ZIKV manipulates host factors to optimize its replication and spread. A key finding of this work is the downregulation of the PS receptors AXL and TIM-1 in ZIKV-infected cells, a previously unappreciated phenomenon. While both AXL and TIM-1 have been previously reported as entry receptors or cofactors to facilitate ZIKV infection (5, 6, 11), we revealed here that their downregulation occurs during the course of infection and is mediated by autophagy and lysosomal degradation. This tactic represents a critical adaptation by the virus to control its infection process, ensuring maximal replication efficiency while minimizing immune activation (**Fig. 5I**). We confirmed that the downregulation of AXL and TIM-1 can occur in all cell types tested, including human epithelial cells, glioblastoma cells, and trophoblasts, underscoring the versatility of this viral infection strategy. This finding is particularly relevant in the context of congenital Zika syndrome, where ZIKV infects early placental and neural tissues, resulting pathogenesis. The downregulation of AXL in trophoblasts may also reduce excessive inflammation in the placenta, protecting fetal tissues from an overactive maternal immune response, while still allowing the virus to establish infection in the developing fetal brain.

We provide evidence that autophagy plays a pivotal role in the downregulation of AXL and TIM-1 process, as inhibiting autophagy with pharmacological agents or genetic knockdowns restored AXL and TIM-1 levels in infected cells; importantly, ZIKV infection itself also induces autophagy. This suggests that ZIKV triggers autophagy as a mechanism to degrade these PS receptors, which is most likely executed by association with the viral proteins NS2A, NS3, and NS4B. The ability of ZIKV to co-opt host autophagy is known to benefit its replication strategy (41), but our findings extend this knowledge by showing that autophagy not only supports viral replication but also modulates the PS receptor availability on the cell surface.

Our findings suggest that the downregulation of AXL and TIM-1 offers several advantages for ZIKV. First, these PS receptors facilitate viral entry, but, once the infection is established, their sustained expression could increase the likelihood of excessive viral entry into infected cells, namely superinfection and immune activation. It’s well established that superinfection can result in viral recombination or cytotoxic effects that would be detrimental to the virus’s ability to replicate efficiently (31, 32). By downregulating these PS receptors, ZIKV minimizes the risk of superinfection, as demonstrated in our dual-infection assays. This would ensure that once a cell is infected, it remains infected at an optimal level, preventing the overburdening of the host cell and potential premature cell death. Moreover, the downregulation of AXL and TIM-1 may aid in immune evasion. These PS receptors are not only viral entry factors but also play roles in modulating innate immune signaling (6, 44). AXL, in particular, has been shown to modulate type I interferon (IFN) responses, which are critical for antiviral defense (45, 46). Our data indicate that ZIKV downregulates AXL to suppress IFN-β and ISG15 expression, thereby dampening the antiviral response of infected cells. This suppression of the type I IFN pathway would allow ZIKV to replicate more effectively within the host by preventing the induction of potent antiviral states in neighboring cells. While we did not find strong effects of TIM-1 in modulating the antiviral type I and pro-inflammatory response in A549 cells, we cannot completely rule out this function in other cell types. More studies are needed. Additionally, the functional consequences of AXL and TIM-1 downregulation extend beyond immune evasion. We observed that ZIKV-infected cells with reduced AXL and TIM-1 expression are less prone to apoptosis, a form of programmed cell death that can limit viral replication by prematurely destroying infected cells. By downregulating these receptors, ZIKV reduces receptor-mediated apoptosis, thus promoting cell survival and allowing for prolonged viral replication. This is particularly important for the virus, as preserving host cell viability is crucial for sustained viral production and spread.

The implications of these findings are broad. They demonstrate that ZIKV has evolved a sophisticated mechanism to balance receptor usage, immune modulation, and cell survival to optimize its infection process. The ability of ZIKV to control receptor levels through autophagy represents a finely tuned strategy that maximizes viral replication while minimizing damage to the host – a process which is likely critical for persistent ZIKV infection (47). However, the role of AXL/TIM-1 downregulation by ZIKV *in vivo* is currently unknown and will be explored in future studies. From a therapeutic standpoint, targeting the autophagy pathway or the specific viral proteins involved in receptor downregulation could provide new strategies for limiting ZIKV replication and spread. Furthermore, our findings raise important questions about the role of receptor regulation in other viral infections, including dengue virus, hepatitis C virus, West Nile virus, and Ebola virus (24, 48). Further studies on PS receptor modulation in other viral infections could provide broader insights into viral pathogenesis and host defense mechanisms.

## MATERIALS AND METHODS

### Cell and Viruses

HEK293T cells, African Green Monkey kidney cells (Vero; ATCC #CCL-81), U87.MG cells, CHME3 cells, and A549 cells were maintained in Dulbecco’s modified Eagle medium (DMEM) supplemented with 10% fetal bovine serum (FBS; Hyclone, logan, UT), 100 U/ml penicillin, 100 μg/ml of streptomycin, and incubated at 37^0^C in 5% CO2. ZIKV strain MR766 (GenBank accession number LC002520), originally isolated in Uganda in 1947, and ZIKV strain FSS13025 (GenBank number: KU955593.1), isolated in 2010 from the blood of a patient from Cambodia, were provided by the University of Texas Medical Branch (UTMB) Arbovirus Reference Collection. Virus stocks were prepared by inoculation onto a confluent monolayer of Vero cells with two rounds of amplification. Routinely the collected ZIKV titer ranged between 1-10 × 10^6^ plaque forming units (PFUs) per ml.

### Plasmids and Reagents

Plasmids WNVII GZ, WNV C, WNV E and WNV M encoding West Nile virus (WNV) genome containing GFP tag, WNV capsid, WNV E protein and WNV M protein respectively were kindly provided by Dr. Theodore C. Pierson(49). ZIKV MR766 E-M harboring conjugated MR766 E and M proteins was synthesized from GenScript. Plasmid Rluc-ZIKV and mCherry-ZIKV were kindly provided by Dr. Pei-Yong Shi at UTMB (50). ZIKV plasmids encoding individual proteins were kindly provided by Dr. Qiyi Tang at Howard University (51).

### Human ESC Culture and Differentiation

The human ESCs (H1, WA01) were cultured and differentiated as previously reported (38). Briefly, cells in six-well tissue culture plates (Thermo Scientific) coated with Matrigel (BD Bioscience) under an atmosphere of 5% (vol/vol) CO_2_/air at 37 °C in mTeSR1 medium (STEMCELL Technologies). Cells were passaged every 5–6 days. For trophoblast differentiation, on the day after passaging onto Matrigel-coated dishes at 1.2 × 10^4^ cells/cm^2^, the culture medium was changed to DME/F12 medium (Thermo Scientific) with knock-out serum replacement (KOSR, Invitrogen) that had been conditioned by mouse embryonic fibroblasts (MEF) and supplemented with FGF2 (4 ng/mL). After 24 h, the conditioned medium was replaced with daily changes of nonconditioned DME/F12/KOSR medium lacking FGF2, but containing BMP4 (10 ng/mL), A83-01 (1 μM), and PD173074 (0.1 μM) (BAP treatment) for 4 days then the cells were subsequently used for ZIKV infection.

### Plaque assay

Infectivity of ZIKV was quantified by plaque assay in Vero-CCL81 cells. Briefly, cells were seeded at 24-well plates at a density of 1 × 10^5^ cells per well. After 24 hours of culture, Vero-CCL81 cell monolayers were infected with 100 μl of a 10-fold dilution series of ZIKV, and the plates were incubated at 37^0^C for 1 hour with gentle swirling every 15 min. After removal of the unbound virus, the cells were overlaid with 0.5 ml of Eagle minimum essential medium (MEM) containing 0.5 % agarose, 5% FBS, 1% sodium bicarbonate, 15 mM HEPES (pH 7.7), and 2 mM L-glutamine. After incubation at 37 ^0^C and 5% CO_2_ for 3 days, the plates were fixed in 3.7% formaldehyde. Plaques were visualized by staining with 0.1% (wt/vol) crystal violet.

### ZIKV reporter virus production

One-round WNV REN reporter virus was generated by co-transfecting plasmids WNVII REN, WNV C, WNV E and WNV M into 293T cells at a ratio of 1:1:0.25:0.25. ZIKV REN reporter virus was generated by co-transfecting plasmids WNVII REN, WNV C, ZIKV MR766 E-M into 293T cells at a ratio of 1:1:0.5. Twenty-four hours after transfection, viruses were collected once every 12 hours.

For replication-competent viruses, i.e., reporter viruses with either mCherry or Rellina luciferase (Rluc) tagged, were generated as previously described (50). Briefly, Plasmid Rluc-ZIKV and mCherry-ZIKV were propagated in *E. coli* Stbl2 and purified using MaxiPrep PLUS (QIAGEN). For *in vitro* transcription, 10 μg of Rluc-ZIKV or mCherry-ZIKV was linearized with restriction enzyme ClaI (NEB). The linearized plasmids were extracted with phenol-chloroform and chloroform, precipitated with ethanol, and resuspended in 15 μl of RNase-free water (Ambion, Austin, TX). The mMESSAGE mMACHINE kit (Ambion) was used to transcribe RNA in a 20-μl reaction with an additional 1 μl of 30 mM GTP solution. The reaction mixture was incubated at 37°C for 2 hr, followed by the addition of DNase I to remove the DNA template. The viral RNA was precipitated with lithium chloride, washed with 70% ethanol, resuspended in RNase-free water, quantitated by spectrophotometry, and stored at −80°C in aliquots. For transfection, approximately 10 μg of RNA was electroporated to 8 × 10^6^ Vero-CCL81 cells in 0.8 ml of Ingenio Electroporation Solution (Mirus, Madison, WI), in 4 mm cuvettes with the GenePulser apparatus (Bio-Rad) at settings of 0.45 kV and 25 μF, pulsing three times, with 3 s intervals. After a 10 min recovery at room temperature, the transfected cells were mixed with media and incubated in a T-175 flask (5% CO_2_ at 37^0^C). At different time points, recombinant viruses in cell culture media were harvested, clarified by centrifugation at 500 × g, stored in aliquots at −80°C, and subjected to analysis.

### Quantification of *Renilla* luciferase reporter viral infection

A549 or Vero cells were seeded on a 12-well plate. Next day, the Rluc reporter viruses were applied to each well at different MOIs. The plates were incubated at 37°C for 1 hour with gentle swirling every 15 min. After removal of the unbound virus, the cells were further incubated at 37°C. At various time points, the cells were washed once with PBS and lysed using cell lysis buffer (Promega). The cells were scraped from plates and stored at −80°C. Once samples for all time points had been collected, luciferase signals were measured in a luminescence microplate reader (BioTek Cytation 5) according to the manufacturer’s protocol.

### Quantification of GFP reporter viral infection

rVSV-G-GFP was applied to pre-seeded A549 or Vero cells in a 12-well plate. Cells were checked once every 2 hours, and then were digested with 0.05% trypsin solution. After fixation under 3.7% formaldehyde, cells were pelleted for flow cytometry analysis (Attune NxT flow cytometer, Invitrogen).

### Generation of knockdown and knockout cells

AXL and TIM-1 genes were knocked out via CRISPR/Cas9 strategy. Briefly, guide RNA targeting AXL gene 5’-CTGCGAAGCCCATAACGCCA-3’, TIM-1 gene 5’-GGACACACGCTATAAGCTAT-3’ and non-targeting oligonucleotide 5’-CACTCACATCGCTACATGA-3’ were introduced into the vector pLentiCRISPR v2 (Genscript). The packaging viruses were produced by co-transfecting 293T cells with 1 µg of pLentiCRISPR v2 based vector, 1 µg pCMVΔR8.2 and 0.5 µg pMDG at ratio of 1:1:0.5. Collected supernatants containing viral particles were applied to A549 cells. Twenty-four hours after transduction, A549 cells were reseeded in DMEM medium containing 5 µg/ml puromycin for one week, allowing for two rounds of selection. Single colonies were selected, and each colony was passaged and genotyped. Initially, the knockdown efficiency of cell culture from each single colony was screened by western blotting. Then the DNA was isolated using a DNeasy Blood & Tissue Kit (Qiagen). The genomic region surrounding the CRISPR/Cas9 target site for each gene was PCR amplified, and PCR products were purified using a QIAquick Gel Extraction Kit (QIAGEN) according to the manufacturer’s protocol. The amplicons were cloned into the pCR-BluntII-TOPO vector (Invitrogen). Each colony was selected, and the amplicon sequences were analyzed using a 3100 Genetic Analyzer (ABI).

Beclin-1 and ATG5 were knocked down in A549 cells by short-harpin (sh) RNAs. Similar to CRISPR/Cas9 procedures, shRNA packaging viruses were produced by co-transfecting 293T cells with 1 µg of lentiviral pLKO.1-puro-based vector (Sigma), 1 µg pCMVΔR8.2 and 0.5 µg pMDG at ratio of 1:1:0.5. A549 cells were transduced with lentiviral shRNA viral particles for 2 hours. Then cells were selected under 1 µg/ml puromycin for 1 week.

### Virus binding assay

To compare the ZIKV binding to Scramble KO and AXL or TIM-1 knockout cells, A549 cells were incubated with viruses containing equal amounts of ZIKV (5 × 10^6^ PFUs) in 1.5 ml Eppendorf tube on ice for 2 h. After the incubation, cells were washed twice with cold PBS, and stained with anti-flavivirus E protein antibody (4G2, Minipore, 1:500) for 1 hours, followed by incubation with secondary antibody FITC-anti-mouse (1:100) for 45 min. All procedures were performed on ice. After fixation under 3.7% formaldehyde, cells were pelleted for flow cytometry analysis (Attune NxT flow cytometer, Invitrogen).

### Immunofluorescence

A549 or embryonic stem cell-derived trophoblast cells infected with ZIKV virus were fixed with 4% paraformaldehyde and permeabilized with permeabilization IC buffer (ThermoFisher Scientific) for 10 min. Cells were blocked with 2.5% BSA/PBS buffer, and stained with anti-Flavivirus E antibody (4G2, Minipore) or anti-AXL antibody (Cat# AF154, R&D) overnight at 4 °C. After 3 washes with PBS, cells were incubated with anti-mouse-TRITC and anti-goat-FITC for 1 h. Cells were stained with DAPI and images were collected using a Leica DMI6000 B inverted deconvolution microscope.

### Western blotting

Western blotting was performed as previously described (52). Briefly, cells were lysed in pre-chilled RIPA buffer (1% NP-40, 50mM Tris-HCl, 150 mM NaCl, 0.1% SDS, and protease inhibitor mixture) or Triton X-100 buffer (1% Triton X-100 in PBS) for 20 min and protein samples were subjected to 10% SDS-PAGE. After proteins were transferred to PVDF membrane, primary antibodies of interest were applied, and protein signals were detected using Amersham Imager 600, GE Health Care. Co-immunoprecipitation assay was performed by using anti-FLAG M2 affinity Gel/beads (Sigma) to incubate with transfected 293T cells lysates (RIPA buffer) overnight at 4°C. Unbound proteins were washed off by 3 times 1 × PBS wash. Samples were finally boiled with 5 × SDS-PAGE loading buffer.

### Assessment of cell viability

A549 viability after ZIKV infection was determined by WST-1 assay as previously described (53). Briefly, 1 × 10^4^ A549 cells were seeded in 96-well plates. ZIKV MR766 was added into each well at an MOI indicated. At 0, 12, 24, 36 and 48 hours after infection, 10 µl of WST-1 substrate was applied to each well, followed by a 15-min incubation at 37°C. After gentle shaking at room temperature for 1 min, absorbance was read at 450 nm (with reference at 620 nm).

### RNA quantification by qPCR

ZIKV-infected A549 cells were collected at 0, 6, 12, 24 hours after infection; total RNA was extracted using RNeasy mini kit (Qiagen). Real-time PCR was conducted by using equal amounts (50 ng) of total cellular RNA using the following primer pairs: Human IFN-β: 5’-AGG ACAGGATGAACT TTGAC, 5’-TGATAGACATTAGCCAGG AG; Human ISG15: 5’-CGCAGATCACCCAGAAGATCG, 5’-TTCGTCGCATTTGTCCACCA; Human CXCL10: 5’-TGGCATTCAAGGAGTACCTC, 5’-TTGTAGCAATGATCTCAACACG; ZIKV-C: 5’ CATCAGGATGGTCTTGGCGATTCTAGC 3’, 5’ CTCGTCTCTTCTTCTCCTTCCTAGCATTGA 3’. The RT-PCR reactions were performed according to the manufacturer’s instruction of 1-step Kit (Power SYBR Green RNA-to-C_T,_ ABI).

### Statistical analysis

All statistical analyses were carried out in GraphPad Prism6, with student t-tests or one-way analysis of variance (ANOVA) used, as appropriate. Data from at least 3 to 5 independent experiments were analyzed.

## ACKNOWLEDGEMENTS

The authors thank Dr. Qiyi Tang from Howard University College of Medicine for providing ZIKV protein constructs. The authors also thank the members of Liu lab and Yu lab for their advice, assistance, and reagents. This work was supported by a fund from The Ohio State University to S.-L.L, and the Major Project of Guangzhou National Laboratory (GZNL2023A01009 and GZNL2023A01005) to J.Y., the National Natural Science Foundation of China (NSFC 82371831 to J.Y.), and the Young Talent Program of China (HJJH22-004 to J.Y.). R.R.M and E.T were supported by NIH R01HD094937.

---

**Figure S1.**
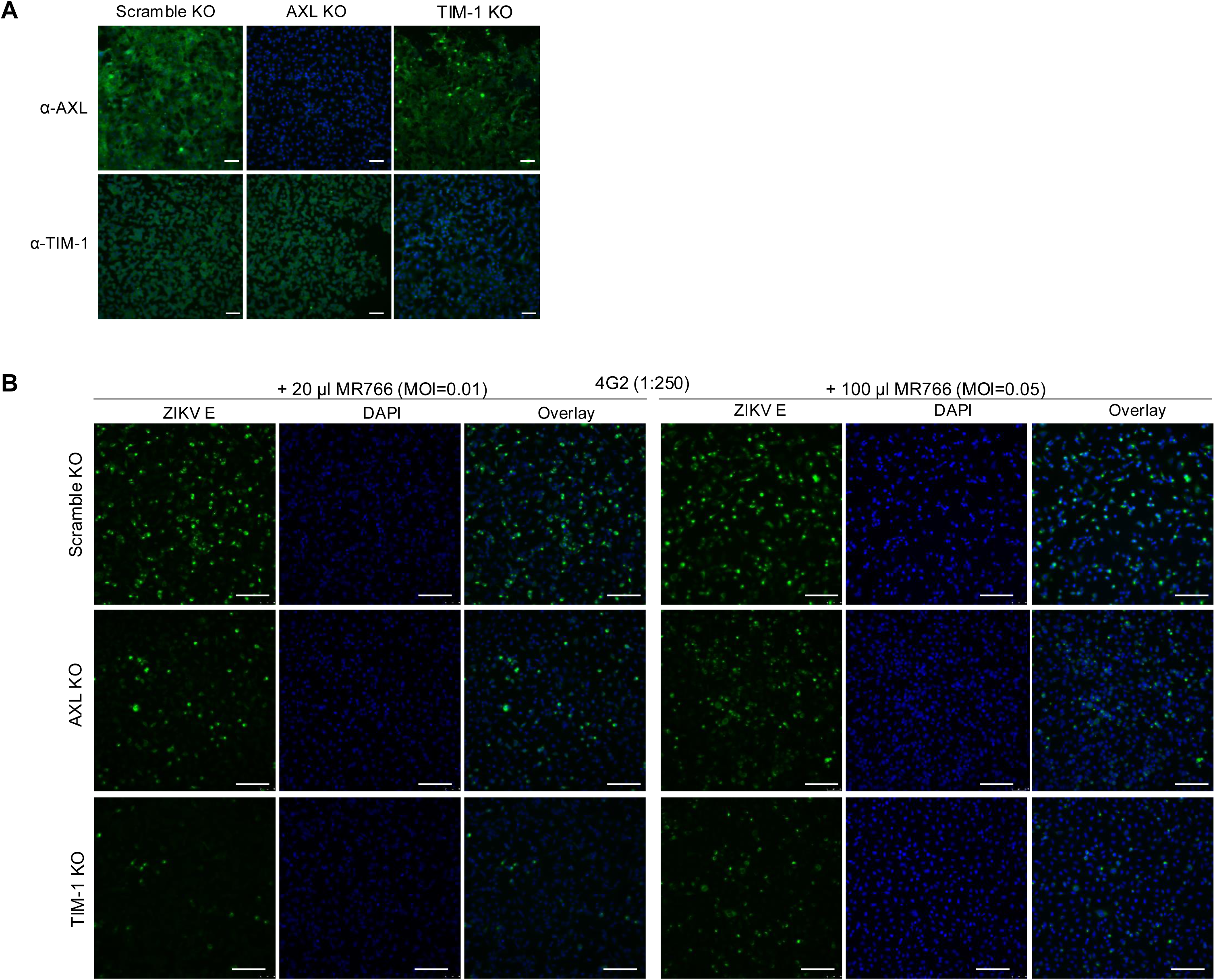
Generation of AXL and TIM-1 knockout A549 cells and their infections by ZIKV. (**A**) The AXL and TIM-1 cell surface expression in was examined by immunofluorescence staining using anti-ZIKV E antibody. Scale bars: 100 μm. (**B**) A549 cells were infected with MR766 at MOI=0.01 or 0.05 for 24 hours, and the infection rate was analyzed by anti-E staining. Scale bars: 100 μm.

**Figure S2.**
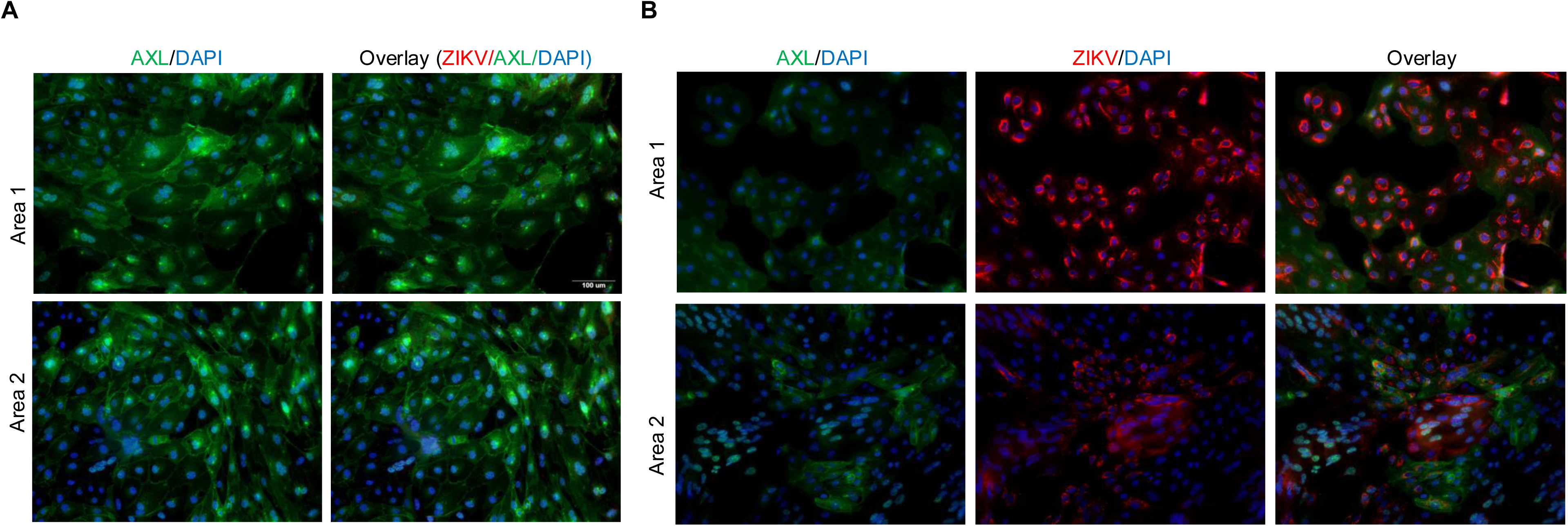
Subcellular localizations of AXL in ZIKV-infected ESCd cells analyzed by immunostaining. ESCd cells were mock infected (**A**) or infected by ZIKV for 36 hours (**B**). Cells were stained by using anti-AXL or ZIKV E protein, followed by fluorescence imaging. Four areas from each slide were randomly selected. Red, ZIKV E; green, AXL; blue, DAPI.

**Figure S3.**
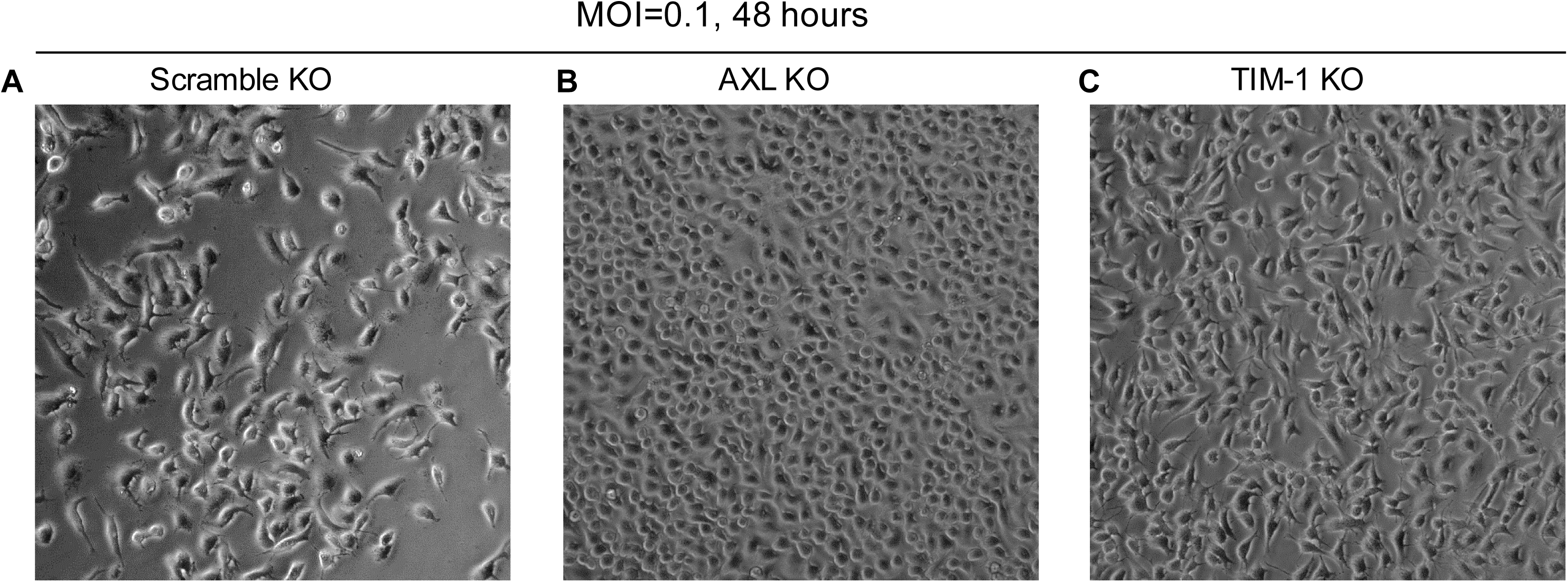
Impact of AXL and TIM-1 KO on ZIKV-induced cell death. **(A)** A549 Scramble KO, **(B)** AXL KO, and **(C)** TIM-1 KO cells were infected with ZIKV MR766 at MOI=0.1 for 48 hours. Cells were imaged after floating cell debris was removed.

